# fuzzyfold: a high-performance framework for stochastic RNA folding kinetics

**DOI:** 10.64898/2026.06.17.732885

**Authors:** Stefan Badelt

## Abstract

The analysis of nucleic acid secondary structures is overwhelmingly dominated by methods that analyze the thermodynamic equilibrium distribution and which ignore all dynamic aspects of nucleic acid folding. Yet, there are numerous popular examples of nucleic acid folding that rely on kinetic models, such as RNA riboswitches or DNA strand displacement systems.

Here, I am presenting `fuzzyfold`, a Rust-based software package for nucleic acid secondary structure analysis with an explicit focus on stochastic modeling. The framework introduces three-way and four-way shift moves with a biophysically motivated rate-model parameterization, and it is developed with an emphasis on both model flexibility and performance, e.g. allowing for the generation of single cotranscriptional trajectories for thousand-nucleotide long RNA molecules in just a few minutes.

The main strength of the `fuzzyfold` package, however, is its focus on user and developer interfaces for long-term development. It provides easily installable command-line interfaces, e.g. for aggregating data from multiple parallel trajectories efficiently into an ensemble-level dynamic analysis. For developers, the code-base supports straight-forward substitution of thermodynamic and kinetic free-energy models, and a flexible library interface with Python bindings, enabling integration of individual components into custom computational workflows.

## 1 Introduction

The nearest-neighbor free energy model for RNA secondary structure prediction has been firmly established for more than three decades, and it remains among the most reliable methods, despite numerous attempts to improve RNA folding predictions with modern machine-learning approaches [16, 37]. Its core strength is the availability of efficient dynamic programming algorithms to calculate a secondary structure equilibrium distribution [28], based on experimentally derived thermodynamic parameters. This allows for calculation of the minimum free energy structure, as well as the probability of observing any particular structure of interest. Over twelve-thousand thermodynamic parameters have been derived for different secondary-structure contexts [41], although the vast majority remain poorly constrained by experiment and can be calculated from a much smaller set of 363 free parameters [1].

Despite the success of thermodynamic predictions, stochastic RNA folding based on nearest-neighbor energies has attracted comparatively little attention. Rather than analyzing the equilibrium distribution, stochastic methods track conformational changes over time. Existing tools include base-pair-level simulators (`Kinfold` [14], `Kfold` [10], `Multistrand` [33]), helix-level approaches (`Kinefold` [44], `RNAkinetics` [8]), and extensions to pseudoknot topologies (`Kinefold`, `StochFold` [38]), each differing in rate model, move set, or scope. Notably, `Kinfold` and `Multistrand` are tightly coupled to specific thermodynamic structure prediction packages – ViennaRNA [27] and NUPACK [9] respectively – which guarantees thermodynamic consistency but limits flexibility for model parameterization.

Researchers have also built deterministic models, for example, [15, 42] partition secondary structures into macrostates and then approximate thermodynamically consistent rates between the macrostates. Alternatively [5, 26] select representative local minimum conformations and estimate transition rates based on free-energy barriers. Importantly, deterministic models give the exact solution to an approximate master equation, and thus are inferior for two reasons: first, they cannot express expected variance from few experimental observations, second, they have a built-in error from the approximation to the true master equation. Also, present deterministic models are tied to a particular macrostate definition, which is based on properties of the energy landscape. This is inconvenient if researchers are interested in the dynamics between specific structures of interest.

Experimental RNA folding kinetics results are typically published at the level of individual examples [6, 17, 23, 45], and it is well-known that nearest-neighbor model stochastic folding can reproduce experimental results based on anecdotal evidence [3, 5, 18, 21, 38]. Yet, the data is not rich enough to extract a set of well-constrained parameters from experiments, because the corresponding simulation results are robust over a range of different parameterizations. For systems where the thermodynamic model is inconsistent with experimental results, it is not possible to identify correct kinetic parameters. However, to develop a better understanding regarding the timescales of RNA folding, it needs not only richer experimental data, but also flexible software capable of testing different combinations of free-energy models, rate models, and move types. The `fuzzyfold` workspace provides a first step in this direction: an initial set of two ViennaRNA-compatible parameter sets [1, 41] with a new tunable three-parameter model for kinetic folding.

This contribution introduces three-way and four-way shift moves, directly inspired by domain-level move types in nucleic acid nanotechnology [4]. These moves allow direct transitions between paired conformations without passing through fully unpaired intermediates, and there is biophysical evidence that three-way shift moves underlie branch migration in three-way junctions [35]. I use an approximate, but more biophysically sound parameterization of shift moves compared to existing approaches [10, 14] and define subsets of shift moves in Sec. 2.3 that satisfy detailed balance and thus could be the basis of refined future rate models.

The core of this work is a high-performance stochastic simulation engine for single secondary structures evolving over time, with efficient parallelization for ensemble dynamics and flexible interfaces for both end users and developers. Currently, `fuzzyfold` provides tools for generating individual stochastic trajectories (Sec. 2.5) and tracing structure ensembles over time (Sec. 2.7), alongside Python bindings that allow rapid prototyping before contributing high-performance Rust code. In Sec. 3.1, I demonstrate the ensemble dynamics of a small, designed RNA, to show how deterministic modeling can give misleading results, even if the structures of interest correspond to local minima in the energy landscape. Finally, I will discuss the impact of three-way and four-way shift moves on dynamics in the context of four-way branch-migration in Sec. 3.2. A detailed, technical protocol using `fuzzyfold` to simulate a natural cotranscriptional system [36] is described in a separate contribution [20].

## 2 Methods

This contribution focuses on the core principles of single-stranded, pseudo-knot free RNA folding kinetics. The ensemble is represented by a thermodynamic energy landscape in terms of states, fitness function, and moves, for a given nucleic acid sequence *σ*. The states correspond to the set of compatible structures Ω_*σ*_. The fitness function is the thermodynamic free-energy evaluation *E*_*i*_ for each structure *i* ∈ Ω_*σ*_. Finally, the moves are reactions (with corresponding reaction rate constants) to define valid transitions between secondary structures. Generally, all moves are treated as “elementary”, which means that they represent direct transitions without metastable intermediates. Stochastic RNA folding is modeled as a continuous-time Markov chain (CTMC), where a state corresponds to a single secondary structure in the thermodynamic ensemble.

### 2.1 State space and energy evaluation

An RNA structure is valid if and only if (i) the base-pairs are *AU, CG* or *GU* (ii) each base forms at most one pair, (iii) base-pairs are nested, (iv) and hairpin-loops have three or more unpaired nucleotides [27, 41]. Then, given a nucleic acid sequence *σ*, the nearest-neighbor energy model assigns a thermodynamic free energy *E*_*i*_ for each compatible structure *i* ∈ Ω_*σ*_. The frequency of observing any secondary structure at equilibrium follows a Boltzmann distribution, hence, the probability of each structure is

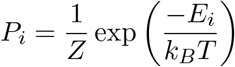

with Boltzmann constant *k*_*B*_, temperature *T*, and partition function 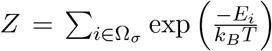. To obey this statistical equilibrium property, the ratio of a rate *k*_*i*→*j*_ and the corresponding reverse rate between any two secondary structures can be constrained by detailed balance

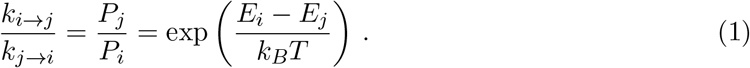

The `fuzzyfold` package supports parameters for RNA and DNA folding. In particular, the RNA(Turner2004) parameters and DNA(RNAstructure) parameters available at NNDB [31, 41], as well as an extended set that includes Pseudo-Uridine [24] and special hairpin parameters [34, 39]. For consistency with thermodynamic modeling, `fuzzyfold` follows the exact implementation of the `ViennaRNA` [27] energy evaluation.

Note that free-energy evaluation is typically constrained to be compatible with efficient dynamic programming algorithms for nucleic acid structure prediction. A popular example is that the free energy of large unpaired loops changes logarithmically with the number of unpaired bases, which is approximated by additive terms in classic nearest-neighbor decomposition schemes. A stochastic simulation is *not* constrained to use an energy model that is suitable for thermodynamic analysis, but it is very convenient if the equilibrium distribution for a system of interest can be calculated directly.

### 2.2 Move sets

Move sets are chosen based on biophysical intuition. A classic choice is the formation and breaking of single base-pairs (see Fig. 1a). Because single base-pair moves are always reversible, reaction rates can be calculated using the detailed balance condition. Furthermore, this move set guarantees that every structure is connected to every other structure in the ensemble through a sequence of moves. Together, these two properties guarantee that the simulated system is ergodic, that is, it reaches the correct equilibrium distribution independently of starting conditions.

**Figure 1.**
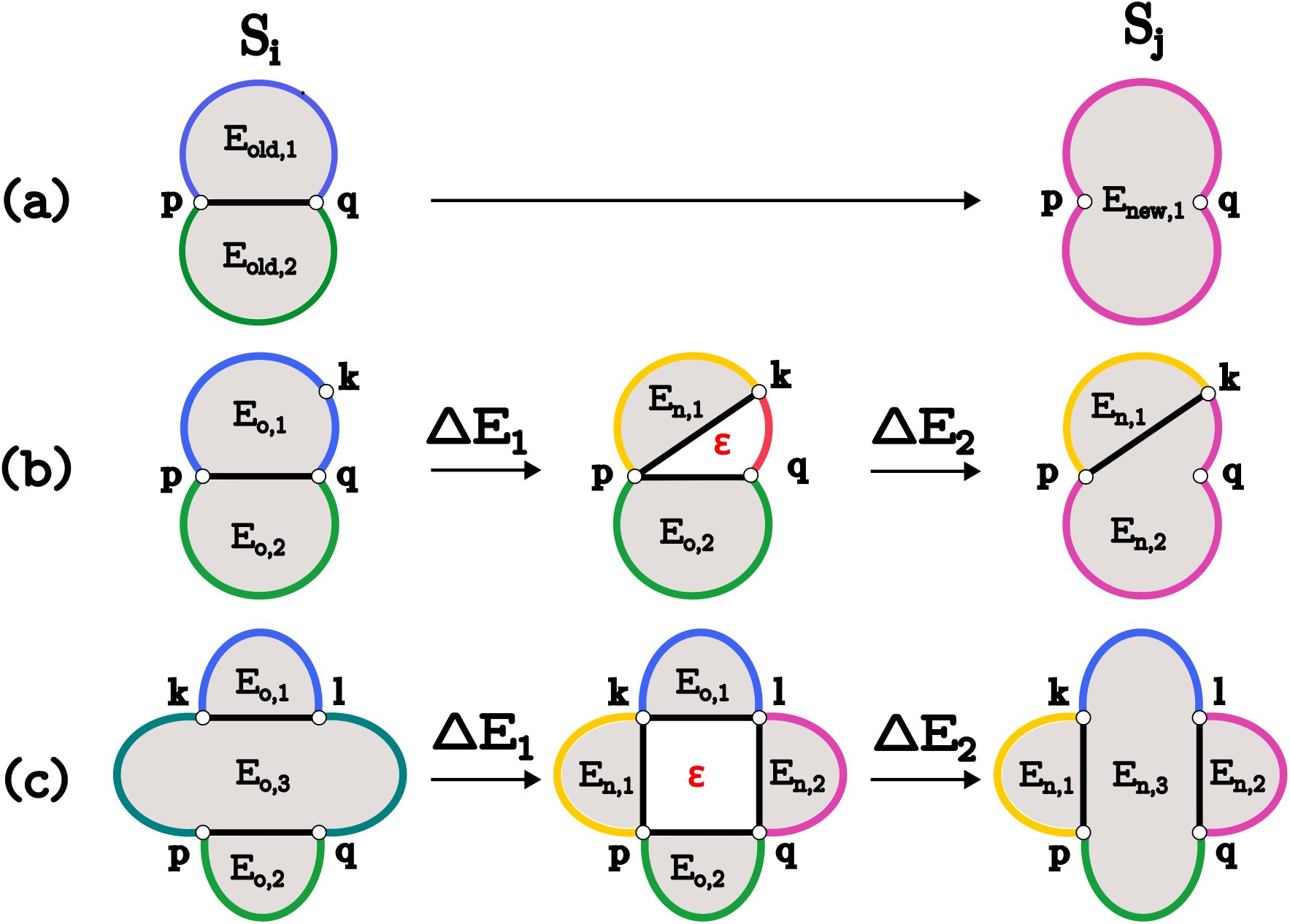
Different types of elementary moves for nucleic acid folding. All moves are reversible, but annotated in forward direction to show the transition from a structure *S*_*i*_ to *S*_*j*_ in terms of the immediately affected loops. Loops are shown with gray background and can have arbitrary type (hairpin, internal, multi, or exterior). **(a)** Deletion of a single base-pair joins two loops *E*_old,1_, *E*_old,2_ into one loop *E*_new,1_. The reverse addition of a base-pair splits one loop into two (not shown). **(b)** A three-way shift move changes a base-pair (*p, q*) directly to some base-pair (*p, k*). The transition is modeled including a transient intermediate forming both base-pairs at once, which involves two standard loops of the nearest-neighbor model, and one unaccounted region *ϵ*. **(c)** A four-way shift move changes two base-pairs (*p, q*), (*k, l*) directly to pairs (*p, k*), (*l, q*). The transient intermediate state forms four base-pairs at once, which involves four standard loops of the nearest-neighbor model, and one unaccounted central region *ϵ*.

Full compatibility with the nearest-neighbor thermodynamic equilibrium distribution is typically violated by simulators using alternative move sets [8, 10, 44]. However, it is possible to include additional moves while maintaining consistency with thermodynamic models. The `fuzzyfold` package supports three-way and four-way shift moves (see Fig. 1), which allow for direct transitions between paired conformations. The former has been known as “shift” moves for RNA folding kinetic simulations [14], and there is evidence that certain conformational rearrangements heavily rely on these types of moves [35]. Since both three-way and four-way shift moves are reversible and any transition achievable by a shift move can equivalently be reached through a sequence of single base-pair moves, the addition of shift moves to an ergodic base-pair open/close system preserves both irreducibility and detailed balance, and therefore ergodicity. This property does not hold for ad-hoc selected subsets of shift moves, unless it can be guaranteed that the moves are reversible. In the context of Fig. 2, I identify subsets of shift moves that are provably reversible, and which may be the basis for refined parameterization in future rate models. Importantly, and in contrast to previous work [10, 14], `fuzzyfold`‘s rate-parameterization for shift moves is based on a biophysically plausible model that estimates the activation energies of different move types (see next section).

**Figure 2.**
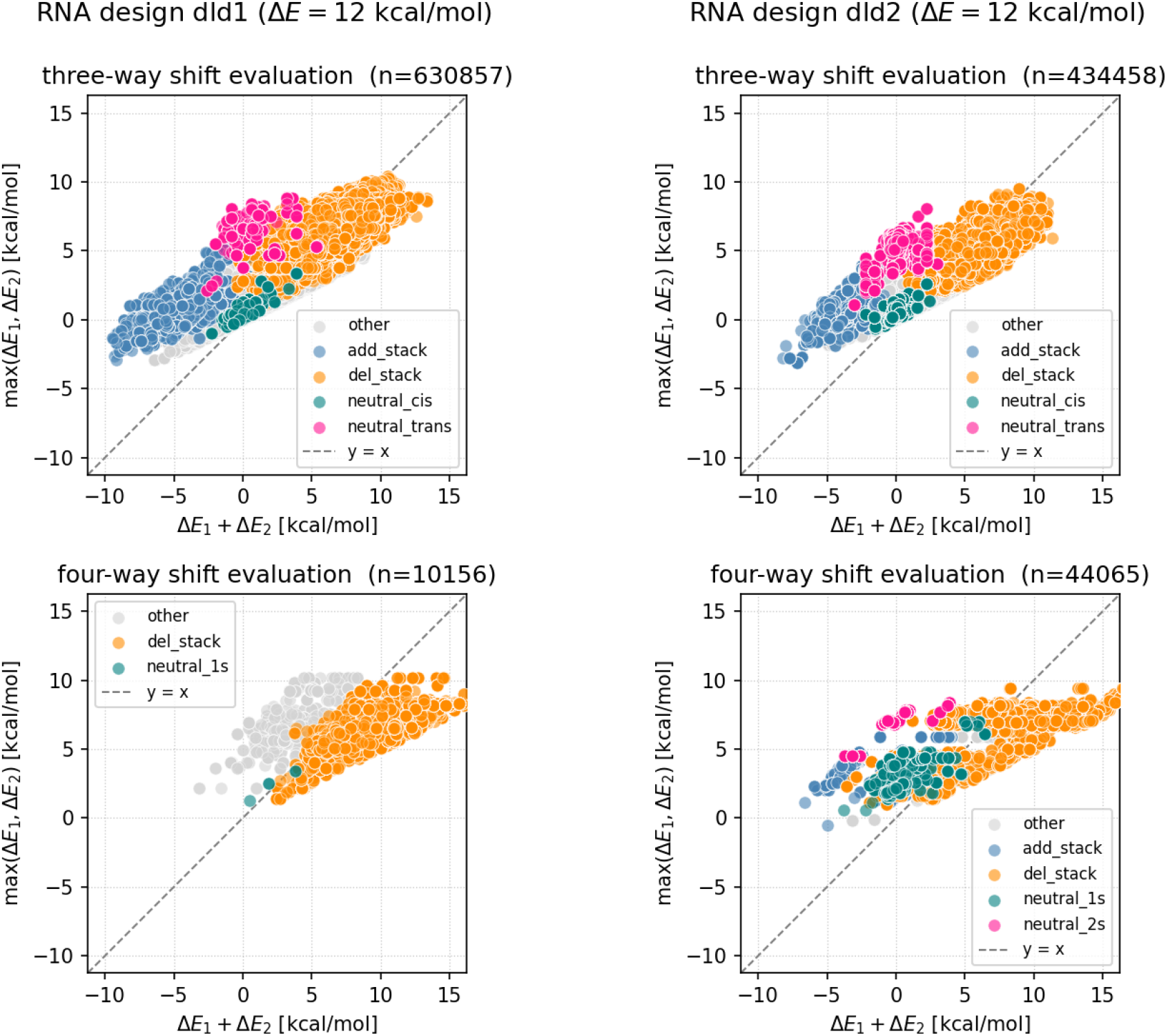
Partial-step barrier max(Δ*E*_1_, Δ*E*_2_) versus overall free-energy difference Δ*E*_1_ + Δ*E*_2_ for shift moves enumerated within 12 kcal*/*mol of the minimum free energy. Each point represents a single shift move. Points above the diagonal indicate moves whose partial-step barrier exceeds the total free-energy difference – these activation energies cannot be captured by a Metropolis-type rate model. Results are shown for two designed RNA sequences (dld1, dld2) described in Secs. 3.1 and 3.2. Move types are classified by their net effect on stacking interactions (see Sec. 2.3).

### 2.3 Rate model

Transition rates follow a standard Arrhenius model with 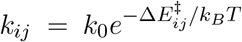, where 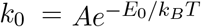 absorbs the attempt frequency A and the reference activation energy *E*_0_ into a single parameter. When only base-pair formation and breaking is allowed, the free-energy difference 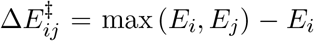 is the activation energy, which is mathematically equivalent to the often used Metropolis model [29] for RNA folding,

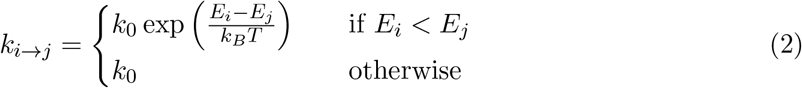

that treats all downhill moves equally fast and scales uphill moves by the full free-energy difference. Throughout this contribution, *k*_0_ denotes the fastest rate for add and delete moves. However, the more general Arrhenius form is necessary to include the activation energies beyond free-energy differences for three-way (*k*_3ws_) and four-way (*k*_4ws_) shift moves. In both cases, the activation energy accounts for an intermediate state where all pairs form at the same time, corresponding to the displacement of present pairs through the formation of invading pairs (see Fig. 1). Formally,

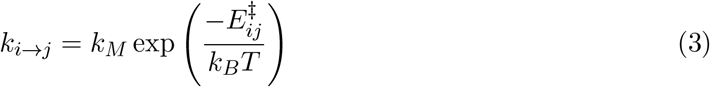

where *k*_*M*_ is the parameter for the fastest moves of type *M* and the activation energy 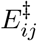 is determined by a two-step picture in which pairs form and dissolve sequentially. In each step, a single nearest-neighbor loop is exchanged, and an additional intermediate loop with unknown free-energy contribution *ϵ* is created or dissolved. Specifically, for a three-way shift move the activation energy is

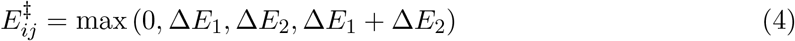

where Δ*E*_1_ = (*E*_new,1_ +*ϵ*)−*E*_old,1_ and Δ*E*_2_ = *E*_new,2_ −(*E*_old,2_ +*ϵ*) are the free-energy costs of the two partial steps. Although *ϵ* cancels in the sum Δ*E*_1_ +Δ*E*_2_, it does not cancel in the individual terms and therefore contributes to the activation energy in general. The contribution of *ϵ* is unknown, and it may be negligible, if the exchange of measured nearest-neighbor loop free-energies sufficiently approximates the intermediate energy barrier. Until experimental evidence suggests otherwise, we assume that *ϵ* is negligible, and allow for calibration to experimental data only by changing the rate parameter *k*_*M*_. The same expression applies to four-way shift moves, with Δ*E*_1_ and Δ*E*_2_ corresponding to the simultaneous formation and dissolution of two pairs respectively (see Fig. 1).

To illustrate the practical implications of this rate model, Fig. 2 shows max(Δ*E*_1_, Δ*E*_2_) against Δ*E*_1_ + Δ*E*_2_ for a variety of three-way and four-way shift moves. The moves were generated from all neighbors of the secondary structures within 12 kcal*/*mol of the minimum free energy, of two designed RNA sequences whose dynamics are discussed in Secs. 3.1 and 3.2. Three-way shift moves are grouped by their effect on stacking interactions: “del-stack” moves lose one stacking pair, “add-stack” moves gain one, “neutral” moves exchange a stack between helices, and “other” collects moves without stacking interactions. As expected, add-stack moves, which are predominantly spontaneous in terms of overall free-energy difference Δ*E*_1_ + Δ*E*_2_, would be assigned a maximum rate *k*_*M*_ under a Metropolis-type model. The transient interrmediate, however, introduces a partial-step barrier as long as one of the two partial steps is uphill. Del-stack moves, by contrast, are reasonably well-described by their overall free-energy difference, with partial-step barriers largely tracking Δ*E*_1_ + Δ*E*_2_.

Surprisingly, there are distinct kinetic clusters among neutral moves that can be cleanly separated into “neutral-cis” and “neutral-trans” subtypes, see Fig. 3. Let (*p, q*) denote the indices of base-pairs such that (*p < q*), then there are four three-way shift moves: (*p, q*) → (*p, k*) is a neutral-trans move, (*p, q*) → (*k, q*) is a neutral-trans move, (*p, q*) → (*k, p*) is a neutral-cis move, (*p, q*) → (*q, k*) is a neutral-cis move. Note that Fig. 3 shows the neutral-trans moves twice, distinguishing the cases where *k* is initially enclosed by, or outside the interval [*p, q*].

**Figure 3.**
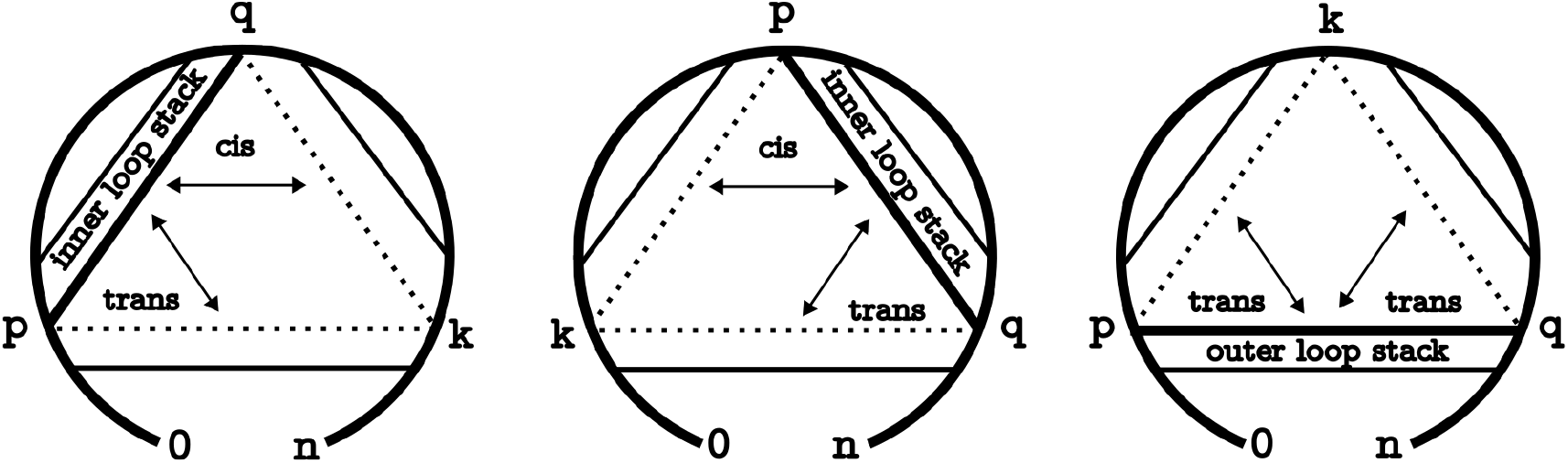
Reversible three-way shift subtypes. The illustration depicts three scenarios with a circular backbone and straight connections for base-pairs. A transition from the pair marked by a bold line to one of the pairs marked by dashed lines presents a three-way shift move. All moves are neutral, in the sense that they preserve exactly one stacking interaction. Different kinetic clusters in Fig. 2 were identified based on the following classification: A trans-move transfers an inner-loop stack (enclosed by the base-pair) to an outer-loop stack (not enclosed by the base-pair) or vice versa. The cis-move also transfers the stack, but it remains the inner loop, that is, both the old and the new pair enclose the stacking interaction.

Loosely speaking, neutral-cis moves are those where new and old pairs are not nested, and thus both pairs (before and after the move) *enclose* a stacking interaction. Those moves tend toward low partial-step barriers and are kinetically fast. In neutral-trans moves, the new and old pair are nested, such that the stacking interaction is *transferred* from the initial pair enclosing a stack to the newly formed pair being the inner pair of the new stacking interaction, or vice versa. Neutral-trans moves exhibit substantially higher barriers. Since the reverse of each subtype is a move of the same subtype, cis and trans represent genuinely distinct reaction classes with different intrinsic timescales. Given the appropriate experimental data, it may be possible to further restrict shift moves based on their subtype. Note, however, that the cis/trans classification depends on the strand indexing convention and may not generalize straightforwardly to multistranded systems where strand order is not uniquely defined.

In four-way shift moves, up to two stacks can be added and deleted at the same time. The moves where two stacks are exchanged at the same time have higher activation energy than those that exchange only one stack. Importantly, however, even in a sequence that was specifically designed to exhibit four-way shift moves, the move type is comparatively rare, and thus accurate parametrization may be more difficult.

### 2.4 Stochastic simulation algorithm

The `fuzzyfold` package implements a Gillespie-type stochastic simulation algorithm [19], where RNA folding is modeled as a continuous-time Markov chain (CTMC) over secondary structures [14]. Each state in the CTMC corresponds to a specific structure *i* with outgoing flux *k*_*i*_ = ∑ _*i* ≠*j*_ *k*_*ij*_, i.e. the sum of reaction rates exiting that structure, where *k*_*ij*_ = 0 if *i* and *j* are not neighbors. The probability of choosing a transition is given by *P*_*ij*_ = *k*_*ij*_*/k*_*i*_ and is famously independent of the waiting time in *i*, which is exponentially distributed with parameter *k*_*i*_ [19].

Per-step runtime depends on the current neighborhood size *N*_*i*_ and the change in neighborhood *N*_*ij*_ = |*N*_*i*_ −*N*_*j*_| induced by each transition. The neighborhood size varies greatly between the open-chain conformation (*N* ≈ *n*^2^) and a fully saturated conformation 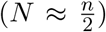. Initial neighborhood calculation is *O*(*N*_*i*_) for both enumerating the neighborhood and calculations of the initial flux. Selection of the next move takes *O*(log(*N*_*i*_)) using a heap [10], updating the neighborhood takes *O*(*N*_*ij*_), and updating the flux takes *O*(*N*_*ij*_ log(*N*_*i*_))) [10]. Consequently, simulations starting in the open-chain conformation are slow, since moves change the neighborhood drastically, whereas simulations in the low energy regime – for example starting from a random Boltzmann-sampled structure – have small, stable neighborhoods and sub-linear per-step costs. Fig. 4 compares the wall-clock runtime of `fuzzyfold` against `Kinfold` [14] of the `ViennaRNA` package [27] and across different simulation modes. The near-linear slopes in the log-log plots reflect the low-energy regime, where per-step costs remain small and stable throughout the simulation. Notably, `fuzzyfold` stores only the current neighborhood, as all attempts at caching or reusing previously computed neighborhoods increased rather than decreased runtime.

**Figure 4.**
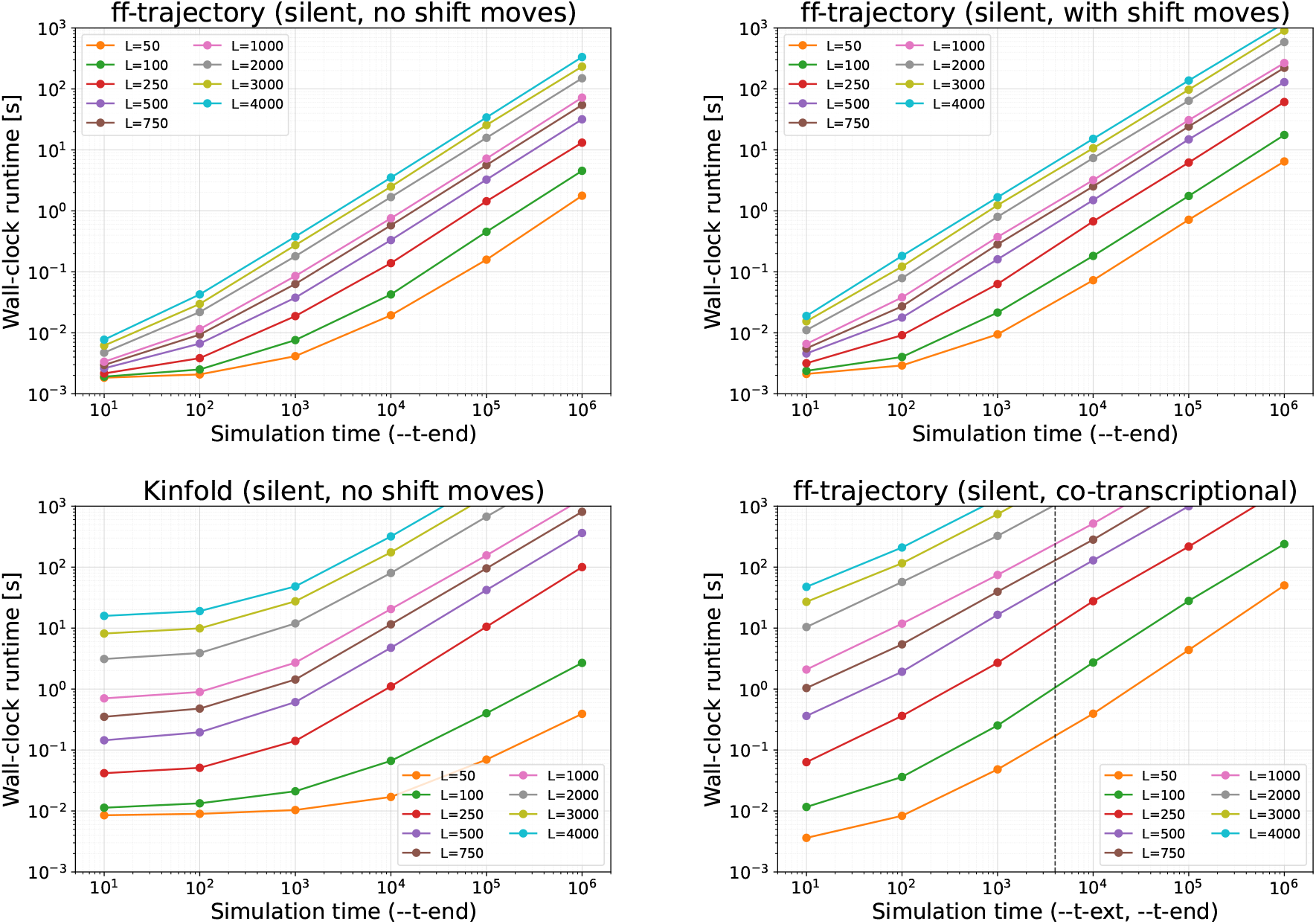
Runtime analysis of single stochastic trajectories using arbitrary time units (*k*_0_ = 1). 100 random sequences were sampled for each length, and then a random (Boltzmann-sampled) starting structure was generated for each sequence. Plots show the median of 100 simulations at fixed simulation times, lines are extrapolations. Typical conversions from the literature suggest that a second in real time is between 10^5^ and 10^6^ arbitrary simulation time units, e.g. simulating a 1000 nt long molecule for one second with `fuzzyfold` takes less than one minute. Note that all simulations use *silent* mode, as runtime increases depending on what information is printed during computation. **(top left)** single trajectories using `ff-trajectory` from `fuzzyfold`. **(top right)** runtime of single trajectories with three-way and four-way shift moves using *k*_0_ = *k*_3ws_ = *k*_4ws_ = 1. **(bottom left)** single trajectories using `Kinfold` from `ViennaRNA`. **(bottom right)** cotranscriptional trajectories using `ff-trajectory`, where the dashed line marks 4000 simulation time units per nucleotide, which is often used in cotranscriptional folding simulations and interpreted as 25 nucleotides per second at *k*_0_ = 10^5^.

### 2.5 Single stochastic trajectories

The program `ff-trajectory` is used to expose single stochastic simulations. While single trajectories are not immediately informative, exposing this interface is important for any customized analysis that does not fit into the pre-built workflows discussed below. It also allows for early access to additional features, before more elaborate interfaces become available. For example, cotranscriptional trajectories have been available already in version 0.4, while cotranscriptional ensemble analysis was only available half a year later in version 0.5. Developers can access simulations of single trajectories through a Python interface, experimenting with alternative workflows before contributing high-performance Rust code for the `fuzzyfold` framework.

### 2.6 From microstates to macrostates

Individual stochastic trajectories (see Sec. 2.5) give results at the level of individual secondary structures, so-called **microstates**. A direct interpretation of results at this level is unpractical, as the number of microstates grows exponentially with sequence length. Instead, a projection to express detailed microstate-level dynamics in terms of a smaller set of distinguishable conformations is necessary. Users define custom **macrostates**, which are non-empty sets of microstates. Macrostates are essential when analyzing the dynamics of RNA molecules with respect to some distinguishable features that may be shared by many structures at the level of microstates.

To help with macrostate definitions, the program `ff-explore` can be used to enumerate macrostates based on two criteria. First, a maximum free-energy difference to a target structure, second, a maximum base-pair distance to the target structure. Both criteria can be used in combination with shift-moves (see Fig. 2), then all structures are enumerated such that the cumulative activation energy for refolding does not exceed the free-energy range. The free-energy criterion is preferred whenever dynamics should be expressed in terms of local minima in an energy landscape. This is very common in RNA research, where landscapes are partitioned into smaller basins to trace their dynamics [15]. However, sometimes a structure of interest is not a local minimum, or part of a shallow basin. In those cases, neighborhood enumeration must be constrained by some distance to the structure of interest.

### 2.7 Stochastic ensemble analysis

The program `ff-timecourse` aggregates data over multiple (parallel) simulations and reports secondary structure occupancies at selected time points. This analysis has the major advantage that the vast majority of output from single trajectories can be discarded on the fly, as only specific time points are of interest. As ensemble diversity at the level of microstates is typically not suitable for distinguishing different functional conformations, `ff-timecourse` provides an interface to define macrostates (see Sec. 2.6) to partition the energy landscape into non-overlapping sets of structures. All structures that are not explicitly listed in any macrostate are collected as “unassigned”, which, itself can be thought of as a macrostate. Automated assignments of structures to macrostates, as well as classifications based on non-exclusive features of secondary structures are ongoing research.

### 2.8 Deterministic ensemble analysis

Various approaches exist to simulate dynamics directly on exhaustively coarse-grained landscapes, [11, 22, 42] or on heuristically coarse-grained landscapes [5, 12, 26]. Coarse-graining first has a notable advantage: ensemble dynamics can be calculated deterministically and over arbitrarily long time periods through matrix exponentiation [42]. While some models are guaranteed to recover the correct equilibrium distribution, all available coarse-grained models introduce biases into the kinetic rates that are difficult to characterize. While `fuzzyfold` does (currently) not support deterministic ensemble analysis, it can be used to evaluate the results of existing deterministic methods. For example, in an often employed deterministic method used in Sec. 3.1, dynamics are expressed in terms of transitions between the lowest gradient-basin macrostates calculated by the software `barriers` [15, 42]. Let *n, δ* and *b* be user-defined parameters, then all secondary structures within some *δ* kcal*/*mol above the minimum free energy are enumerated [43], and assigned to the lowest of *n* reachable minima with basin-height of at least *b* kcal*/*mol. Note that this macrostate definition is based on properties of the landscape, and may not be suitable for tracing the dynamics of specific structures of interest. The “fifth” local minimum in Fig. 5, must be included, and the vast majority of structures with no similarity to the macrostate representatives are distributed among the six macrostates.

**Figure 5.**
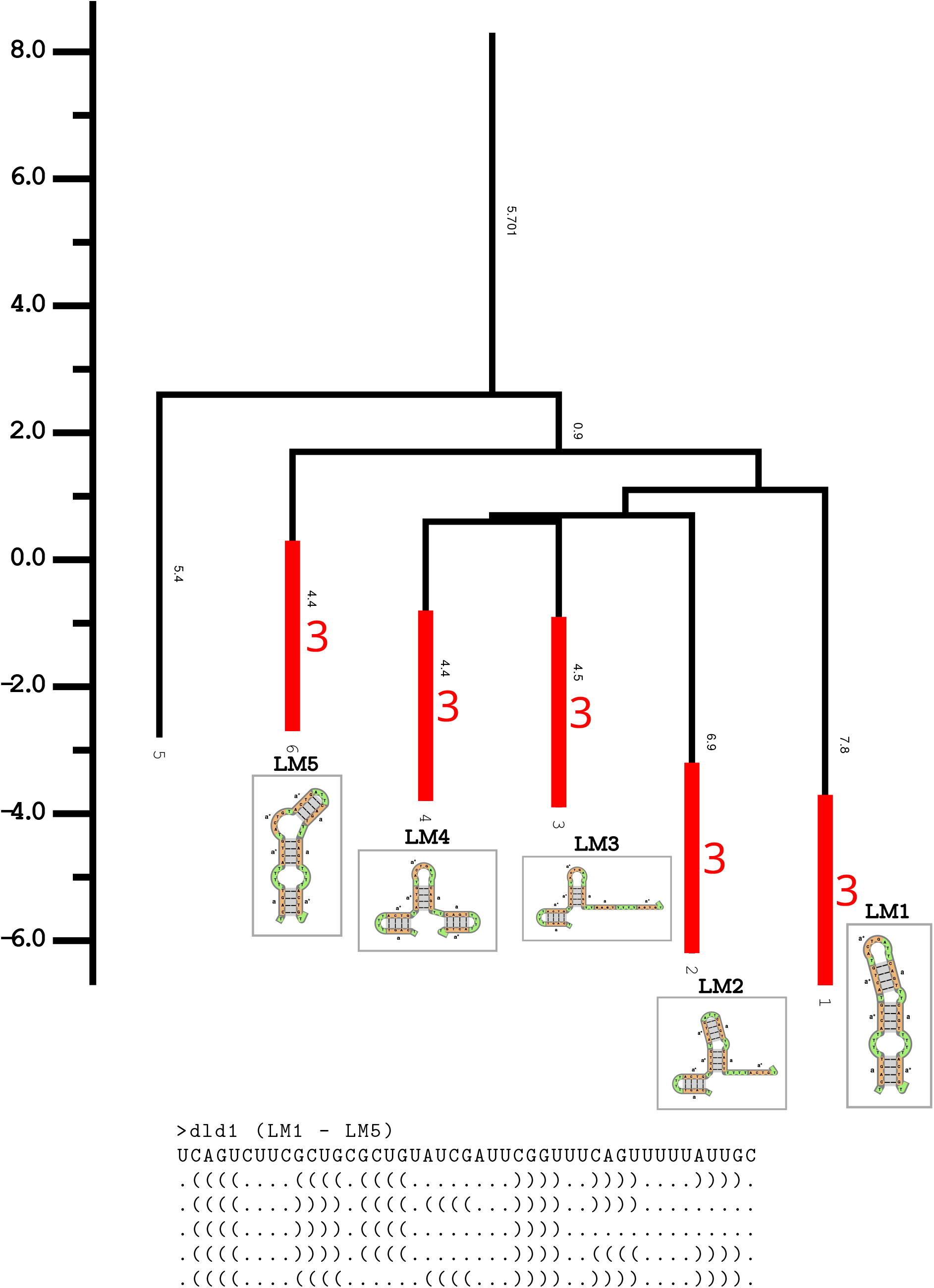
Free-energy landscape of a short, multi-stable, sequence. **(top)** A barrier tree produced by the program `barriers` expressing the free-energy landscape in terms of the free energies of local minima. Suboptimal structures in an energy range of 15 kcal*/*mol were partitioned into *n* = 6 basins of minimal height *b* ≥ 4. Black numbers denote the vertical branch lengths, and thus the actual basin-depth in kcal*/*mol. Basins annotated with structure images, 1, 2, 3, 4, 6, correspond to named structures of interest (LM1 - LM5). The red annotations mark basin heights of 3 kcal*/*mol that were used for custom17 macrostate definitions in `fuzzyfold` analysis. **(bottom)** Sequence and secondary structures of interest in order (LM1 - LM5).

## 3 Results

Two case studies are presented based on designed molecules. First, I am comparing the detailed, stochastic ensemble dynamics (see Sec. 2.7) with coarse-grained, deterministic ensemble dynamics (see Sec. 2.8). Second, I compare the effect of different move types and associated rate constants on the dynamics of a single branch-migration reaction.

### 3.1 Detailed vs coarse-grained dynamics

An existing method for coarse-grained analysis based on the program `barriers` (see Sec. 2.8) is applied to an RNA “dld1” (for domain-level design 1) with five structures of interest (see Fig. 5). All secondary structures within 15 kcal*/*mol above the minimum free energy are enumerated, and assigned to the lowest 6 **local minima (LM)** with basin-height of at least 4 kcal*/*mol. This works well for a designed molecule, because the structures of interest do indeed correspond to local minima in the free-energy landscape. In comparison, `ff-timecourse` is used to simulate dynamics at the detailed level of microstates and then results are expressed in terms of custom-defined macrostates. Both models use the same add and delete moves at *k*_0_ = 10^5^, and no shift moves. Macrostates are defined by enumerating each target structure’s basins up to 3 kcal*/*mol (see Fig. 5). The remaining conformations remain “unassigned”, e.g. the “fifth” local minimum in Fig. 5 is not included. Here, the choice of 3 kcal*/*mol is a compromise between allowing for alternative conformations, and ensuring that a few base-pairs remain intact in all target helices. The conceptually relevant distinction is between structures of interest and the bulk of unassigned structures. The gradient-basin model cannot make this distinction by construction, because every microstate is assigned to one of the leaves of the barrier tree.

As can be seen in Fig. 6, the approximate dynamics using the deterministic, coarse-grained approach can be quite different from the underlying detailed stochastic model. Aggregating over 100-thousand simulations may seem excessive, but ensures that the stochastic sample variance is negligible. First, the time-scales between basin transitions differ from detailed simulations by orders of magnitude. This is expected, because coarse-grained simulations assume instant equilibration within basins, and, in some cases, this can be corrected with a constant factor. Second, gradient-basin assignments can attribute structures to macrostates even though they are structurally unrelated to that macrostate – an artifact that is visible at equilibrium in all simulations, where a substantial fraction of the unassigned ensemble is attributed to LM1. The magnitude of this effect depends on the macrostate definition, but with the present assignment it is particularly pronounced in the simulation starting in LM5, where almost all transition structures are attributed to LM1. Third, simulations from LM4 show that the probability of observing LM3 in detailed stochastic simulations is higher than estimated by deterministic gradient-basin dynamics. Considering, that the gradient-basin LM3 is larger than macrostate LM3 in the stochastic model, this suggests that the deterministic model does not capture the correct detailed transitions between LM4 and lower-energy minima.

**Figure 6.**
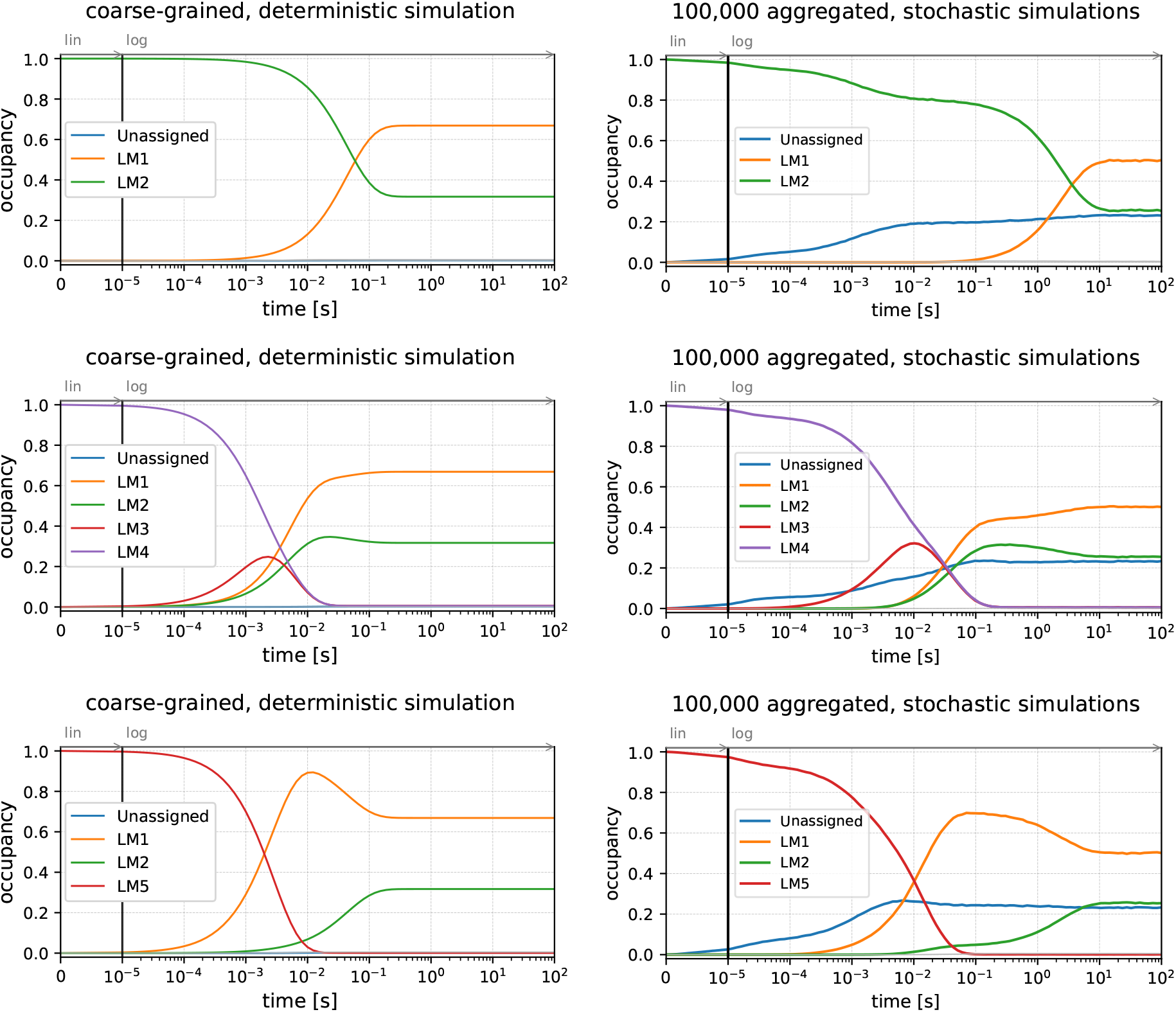
Analysis of ensemble dynamics. Conversions to seconds use *k*_0_ = 10^5^ s^−1^. **(left)** Approximate, deterministic simulations using gradient-basin macrostates, each panel starting from a different local minimum conformation. **(right)** Aggregated data from detailed stochastic simulations mapped into custom-defined macrostates and a remaining set of unassigned structures.

### 3.2 Effect of move type and rate parameters

The *k*_0_ parameter for kinetic RNA folding is a crude approximation to relate RNA folding simulations to real time. Early experiments suggested that base-pair formation is on the order of 10^6^ s^−1^ to 10^7^ s^−1^ [30]. This is consistent with more recent estimates to simulate experimentally tractable structure formation for both RNA and DNA based experiments [13, 40, 46, 47]. In cotranscriptional folding settings, where the folding time per nucleotide must be consistent with experimentally confirmed transcription rates, comparisons of base-pair level simulations with experimental data often use *k*_0_ = 10^5^ s^−1^ [2, 5, 21, 32], which is also used consistently throughout this manuscript.

Here, an RNA is designed to rearrange dynamically via four-way branch migration, and it is investigated how different combinations of parameters influence folding kinetics. All simulations start in structure LM4 (see Fig. 7), and thus should capture the same random walk process in which LM1 and LM3 are visited first at the approximately same time, before the system transitions into its equilibrium distribution. The first simulation uses *k*_0_ = 10^5^ s^−1^, and *k*_3ws_ = *k*_4ws_ = 0 s^−1^, where LM3 and LM1 reach their maximum occupancy after 1 and 10 seconds respectively. Changing three-way-shift moves to *k*_3ws_ = 10^4^ s^−1^ speeds the transitions up by approximately one order of magnitude. Interestingly, additionally allowing four-way-shift moves with *k*_4ws_ = 10^4^ s^−1^ does not seem to have an observable effect on dynamics, suggesting that four-way branch-migration reactions may actually rearrange predominantly using three-way-shift moves.

**Figure 7.**
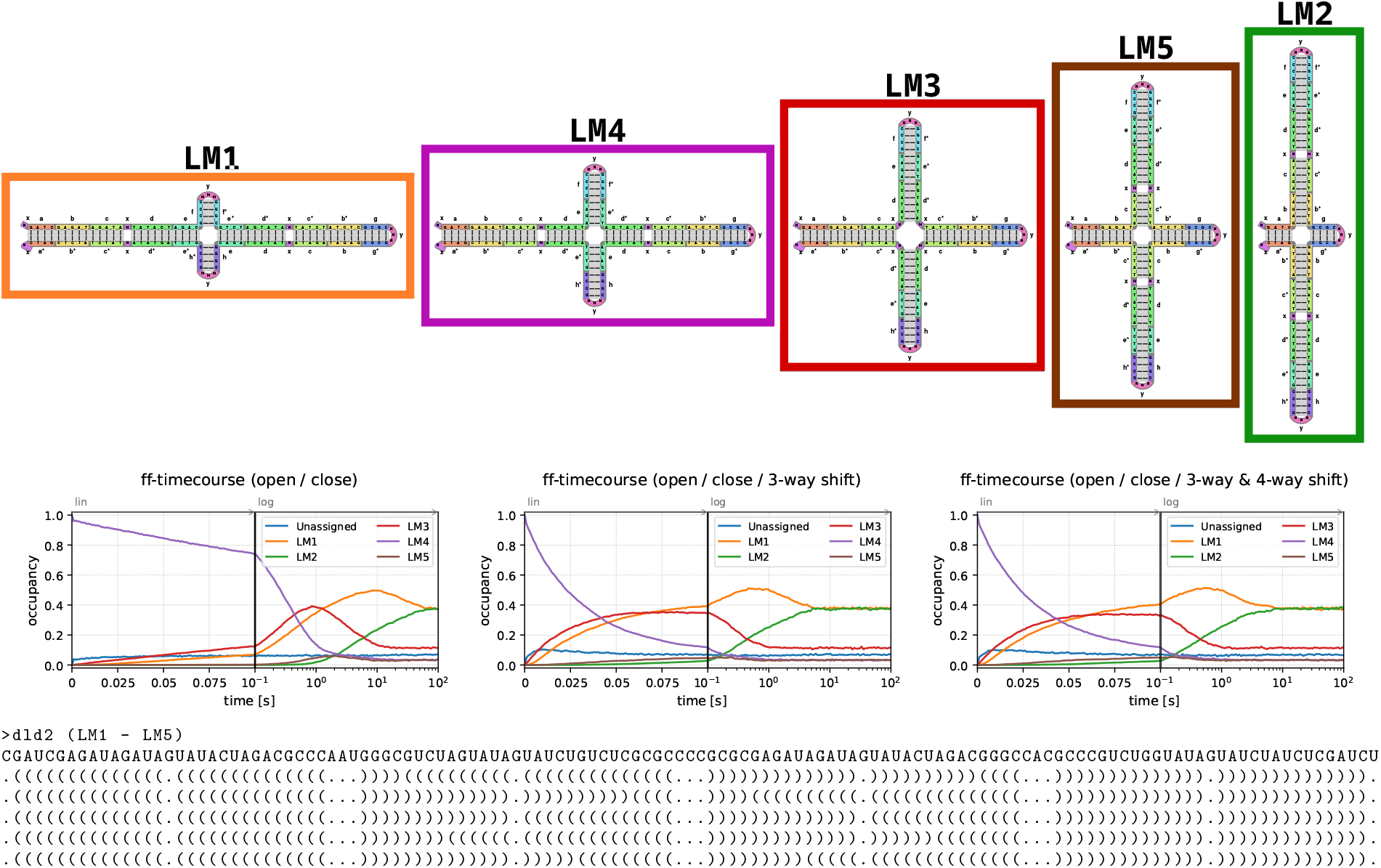
Dynamics of single-stranded four-way branch-migration rearrangements. **(top)**: five different configurations of an RNA molecule. LM1 and LM2 are the lowest free-energy conformations. Colors correspond to `ff-timecourse` results. **(middle)**: Three ensemble analyses of 10^4^ detailed simulations starting in LM4, all using base-pair add and delete moves with *k*_0_ = 10^5^ s^−1^. The first simulation uses only base-pair opening and closing, the second uses additional three-way-shift moves *k*_3ws_ = 10^4^ s^−1^, the third uses three and four-way-shift moves *k*_3ws_ = *k*_4ws_ = 10^4^ s^−1^. **(bottom)** Sequence and secondary structures of interest in order (LM1-LM5).

## 4 Discussion

Among existing kinetic folding tools, most are either tightly coupled to specific thermodynamic packages or lack the flexibility to test alternative rate models and move sets. The `fuzzyfold` package addresses both limitations by separating the simulation engine from the analysis layer, and by exposing the core functionality through both high-performance Rust interfaces and Python bindings for rapid prototyping. The separation of trajectory and ensemble analysis reflects a deliberate choice: single trajectories offer maximum flexibility for custom analysis, while ensemble-level aggregation via `ff-timecourse` provides an efficient path to occupancy dynamics without storing full trajectory data.

Runtime analysis in Fig. 4 between `Kinfold` and `ff-trajectory` shows a significant initial cost in `Kinfold` simulations. This is largely because `Kinfold` computes the minimum free energy (MFE) structure at the start of the first simulation. For short sequences (less than 100 nucleotides) this makes almost no difference, but for very long sequences, it emphasizes a regime in which stochastic simulations with runtime *O*(*N*_*ij*_ log(*N*_*i*_)) per step (see Sec. 2.4) can be much faster than *O*(*n*^3^) MFE computations. Runtime comparisons with other simulation packages were avoided, as runtime depends also on the choice of moves and the kinetic model. For example, simulations that equilibrate faster have reduced runtime, even if they simulate over the same time period. That is, because the neighborhoods of low-energy conformations are smaller. For example, Kfold uses the Kawasaki rate model [25], which is designed to achieve fast equilibration, rather than biophysically correct dynamics. Also, the description of the move-set in [10] is insufficient to verify whether it satisfies detailed balance.

Cotranscriptional trajectories are now much faster than in `Kinfold`. A cotranscriptional trajectory of a 750 nucleotide long random sequence with typical 4000 arbitrary time units per nucleotide [3, 5, 21], takes about 2 minutes on a single core. The runtime for a single trajectory is critical, as this is the smallest unit that can be efficiently parallelized. 100 trajectories already give a good estimate for expected dynamics and can easily be computed in parallel on modern compute architectures. Using `ff-timecourse`, cotranscriptional dynamics can then be traced with respect to specific macrostates of interest. By comparison, the program `DrTransformer` [5], a heuristic cotranscriptional analysis has to identify macrostates during the simulations, and thus takes multiple days for an RNA of the same length.

The case study in Sec. 3.1 demonstrates that detailed stochastic ensemble dynamics can differ substantially from coarse-grained deterministic approximations. The `fuzzyfold` package now provides a computational framework to guide more systematic parameter exploration, in which aggregated stochastic simulations can be directly compared against experimental observables across a range of nucleic acid systems. The new formulation of shift moves and their geometric subtypes offers a flexible basis for building more robust biophysical rate models applicable to a diverse set of rearrangements, potentially refined by secondary-structure context [47]. DNA nanotechnology offers a particularly rich source of experimental constraints through strand displacement systems, which involve the same types of base-pair rearrangements studied here, and branch migration in single-stranded RNA cotranscriptional folding [7] suggests that those parameterizations can be used across different nucleic acid systems. Perhaps disappointingly, the results also suggest that four-way shift moves (and also higher-order shift moves) may not be very useful for a better kinetic model after all.

Finally, the improvement of runtime performance over existing software may also stimulate entirely new research directions. For example, it may cause a paradigm shift in sequence design, where focus has been exclusively on thermodynamic methods, arguing that kinetic simulations are too slow. With sequence design pushing into the regime of long sequences, such as mRNAs for biotechnological applications, kinetic approaches may be the basis of more performant design approaches.

## 5 Conclusion

The `fuzzyfold` package was designed from the ground up with kinetic modeling as its primary objective, rather than as an extension of thermodynamic analysis tools. It provides a modern, flexible, and efficient open-source framework for stochastic RNA folding, with command-line tools for end users, a library interface for trajectory simulations and ensemble analysis, and Python bindings for custom workflows. The choice of Rust facilitates high-performance implementations with straightforward installation and reproducible builds.

As a pure Rust workspace, `fuzzyfold` is straightforward to install and extend, and additional functionality can be contributed as independent crates without modifying the core engine. Possible extensions include multistranded interactions, circular molecules, pseudoknotted conformations, DNA/RNA hybrid systems, and modified nucleotides. Unlike thermodynamic methods, stochastic simulations are not constrained to energy models compatible with efficient dynamic programming, leaving room for future extensions to pseudoknots, non-local free-energy effects, and context-dependent rate parameters.

## 6 Acknowledgements

I want to thank Ivo L. Hofacker for discussions on stochastic simulation performance using log(*n*) flux updates, as well as for helpful discussions on rate-models for three-way shift moves. I want to thank Hannah Hartung, Daniel Guerguerian, and Jonas Jäger for their ongoing efforts to integrate new features into the fuzzyfold workspace, and Hannah Hartung and Daniel Guerguerian additionally for their contributions to the protocol paper accompanying this work. I want to thank Ronny Lorenz for help with ViennaRNA compatible free-energy evaluations, and Leonhard Sidl for critical feedback on the manuscript.

## Notes

https://github.com/bad-ants-fleet/fuzzyfold

### Competing Interest Statement

The authors have declared no competing interest.

### Summary of Updates

Revised introduction and discussion, as well as the justification for the shift rate model. A new Figure 3 shows the difference between neutral-cis and neutral-trans three-way shift moves.

